# Light-inducible generation of membrane curvature in live cells with engineered BAR domain proteins

**DOI:** 10.1101/2020.02.20.958611

**Authors:** Taylor Jones, Aofei Liu, Bianxiao Cui

## Abstract

Nanoscale membrane curvature is now understood to play an active role in essential cellular processes such as endocytosis, exocytosis and actin dynamics. Previous studies have shown that membrane curvatures directly affect protein functions and intracellular signaling. However, few methods are able to precisely manipulate membrane curvature in live cells. Here, we report the development of a new method of generating nanoscale membrane curvature in live cells that is controllable, reversible, and capable of precise spatial and temporal manipulation. For this purpose, we make use of BAR domain proteins, a family of well-studied membrane-remodeling and membrane-sculpting proteins. Specifically, we engineered two optogenetic systems, opto-FBAR and opto-IBAR, that allow light-inducible formation of positive and negative membrane curvature respectively. Using opto-FBAR, blue light activation results in the formation of tubular membrane invaginations (positive curvature), controllable down to the subcellular level. Using opto-IBAR, blue light illumination results in the formation of membrane protrusions or filopodia (negative curvature). These systems present a novel approach for light-inducible manipulation of nanoscale membrane curvature in live cells.

**Highlights:**

- Opto-FBAR enables light-inducible positive membrane curvature formation.
- Opto-IBAR enables light-inducible negative membrane curvature formation.
- Light-inducible activation enables precise spatial and temporal control.
- Opto-BAR systems present a new approach for studying membranes in live cells.

## Introduction

Nanoscale (<1 *μ*m radius) membrane curvature is believed to actively participate in a diverse set of intracellular processes such as endocytosis, exocytosis, and cell migration.^1^ The modulation of nanoscale membrane curvature has been proposed to be an important regulator of cell membrane biology. Membrane curvature generation can induce both physical and biochemical responses from proteins via alteration of the spatial arrangement and binding affinity of membrane-associated proteins as well as stimulation of enzymatic function, respectively.^2,3^ Dysregulation of membrane curvature is often observed in metastatic cancer cells, which exhibit high densities of membrane protrusions and can form tunneling nanotubes to facilitate intercellular communication and transport.^4,5^ Given the numerous observations of relationships between membrane curvature and protein activity, as well as the fact that membrane curvature is regulated by proteins in both the intracellular and extracellular spaces,^6,7^ it has been suggested that the generation of nanoscale membrane curvature itself can induce intracellular signaling responses.^8^ We recently found that nanoscale membrane curvatures underlie how substrate topography alters intracellular signaling, referred to as the “curvature hypothesis.”^8,9^

To date, controlled generation of nanoscale membrane curvatures and their effects on protein function have mostly consisted of *in vitro* studies with liposomes or supported bilayers.^10,11^ *In vitro* findings require validation in live cells, where methods to control the generation of membrane curvature are limited. Vertical nanostructures have been demonstrated as a platform to induce precisely controlled curvatures on the cell membrane, which has been successfully used to probe curvature-dependent endocytosis^12^ and actin polymerization.^13^ But as this form of curvature generation is static, observation of the cell’s response to membrane curvature generation in real time remains difficult. To address these limitations, we sought a new method of generating nanoscale membrane curvature that is dynamic, reversible, and capable of stimulating a large population of cells.

Optogenetics has greatly advanced our understanding of intracellular processes and signaling.^14^ By coupling protein function to a light stimulus, one can study the cellular response to that function with precise temporal and spatial control. A recent study shows that light-induced crowding of a nonfunctional fluorescent protein on the membrane causes membrane deformation, but the degree of deformation and the sign of the induced membrane deformation is not controllable.^15^ We hypothesize that an optogenetic approach of coupling a blue light photoreceptor to proteins capable of generating membrane curvature could provide a means to control nanoscale membrane curvature with blue light.

Among the well-studied classes of proteins capable of generating membrane curvature are members of the Bin/Amphiphysin/Rvs (BAR) domain superfamily. BAR domains consist of coiled-coil dimers that bind to membranes, often with a surface of positively charged amino acids.^16^ Members are further classified into subfamilies based upon the characteristics of their BAR domain. The F-BAR (extended Fes-CIP4 homology (EFC)/FCH-BAR) domain subfamily of proteins are banana-shaped with positive charges on the concave side, enabling them to sense and induce positive curvature (bending membranes toward the protein binding leaflet, Figure 1A, B). I-BAR (IRSp53-MIM homology domain I-BAR/inverse-BAR) domain proteins, conversely, are cigar-shaped with positive charges on a convex surface and can generate negative membrane curvature (bending away from the binding leaflet, Figure 1C).^17^ Both of these families exhibit the ability to oligomerize at the plasma membrane in a concentration-dependent manner, providing a scaffold that promotes bending of the plasma membrane towards their respective curvature preference.^18^

**Figure 1:**
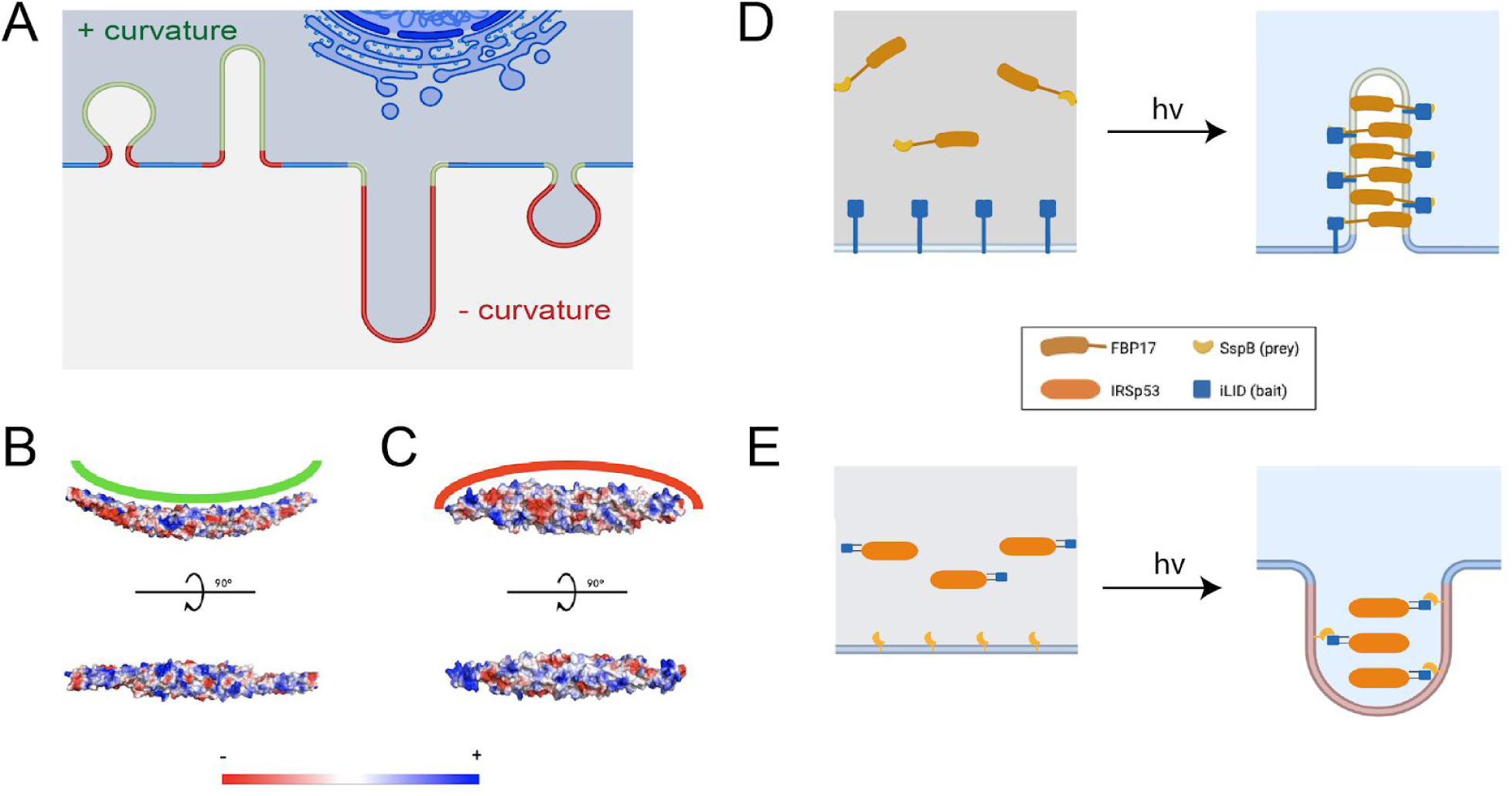
Design principles of opto-FBAR and opto-IBAR. **A**) Schematic depicting positive (green) and negative (red) curvatures of the cell membrane. **B**) Electrostatic surface potential of the FBP17 FBAR domain (PDB ID: 2EFL). **C**) Electrostatic surface potential of the IRSp53 IBAR domain (PDB ID: 1Y2O). **D**) Schematics of opto-FBAR. Blue light illumination recruits the cytosolic FBAR protein to the plasma membrane through the light-gated prey-bait interaction. High concentration of FBAR proteins will self-assemble and generate positive curvature on the plasma membrane. **E**) Schematics of opto-IBAR. Blue light illumination results in the membrane recruitment of the cytosolic IBAR protein and the generation of negative curvature.

Here, we report the development of two optogenetic systems: opto-FBAR and opto-IBAR. Derived from the human FBP17 and IRSp53 proteins, respectively, these systems allow us to induce the formation of positive and negative membrane curvature with fine spatiotemporal precision. Both of these systems utilize the improved light-inducible dimer system (iLID) to force recruitment of the BAR domain protein to the plasma membrane, increasing their local concentrations to promote their oligomerization and induction of curvature.^19^ In the case of opto-FBAR, blue light activation results in the formation of tubular membrane invaginations, controllable down to the subcellular level. For opto-IBAR, in addition in iLID, we incorporated a second photosensor pdDronpa to cage IBAR activity in the absence of blue light. Opto-IBAR activation results in the formation or elongation of cellular membrane protrusions and filopodia.

## Results and Discussion

### Principles and design of opto-FBAR and opto-IBAR

BAR domains are known to assemble on the lipid membrane and form membrane tubules of high curvatures on liposomes *in vitro* in a concentration-dependent manner.^20^ This concentration dependence provides a means by which curvature induction can be modulated using light-induced protein translocation systems. These protein translocation systems consists of two components, hereafter referred to as “bait” and “prey.” In the case of optogenetic translocation systems, the bait protein’s ability to bind to the prey depends on a light stimulus. Upon light activation, the bait and prey dimerize, which can be used to modulate the subcellular distribution of a protein of interest when one component is targeted to a specific region with a signaling peptide.^21^ We hypothesize that by genetically fusing one component of an optogenetic dimerization system to a curvature-inducing protein such as a BAR domain and the other component to a plasma membrane targeting sequence, we can dynamically inducing membrane curvatures with blue light (Figure 1D,E).

To engineer an optogenetic system for positive membrane curvature, we chose formin-binding protein 17 (FBP17). FBP17 is one of the most extensively studied FBAR proteins, known to play roles in a variety of biological processes including clathrin-dependent and -independent endocytosis, cell migration, phagocytosis, and cancer cell invasion.^11,22–25^ Crystal structures of FBP17 are available for the FBAR domain in both polymerized and unpolymerized forms.^26,27^ In a similar vein, to engineer a system for light-induced negative membrane curvature, we chose to engineer the IBAR domain containing protein Insulin Receptor Substrate protein of 53 kDa (IRSp53), one of the more thoroughly characterized IBAR domain containing proteins. IRSp53 has been shown to induce negative membrane curvature via polymerization of its IBAR domain.^28^ The structure of IRSp53’s IBAR domain has been solved and the IBAR has been shown *in vitro* to exhibit similar behavior to other BAR domain proteins, albeit sensing/generating negative membrane curvature.^29,30^

We chose the iLID/SspB optogenetic dimerization pair to engineer our opto-BAR systems (Figure 1D,E). iLID, engineered by photocaging the SsrA peptide with the *Avena sativa* phototropin 1 LOV2 domain, binds to SspB upon blue light illumination with fast on/off kinetics.^19^ The group that engineered iLID also generated multiple versions of SspB, dubbed nano, micro, and milli, with different affinities for the SsrA peptide.^31^ These varieties make it possible to fine tune the kinetics and affinity if necessary.

### Construction and optimization of the opto-FBAR systems

For our initial investigation, the prey component of the optogenetic FBAR constructs (Figure 2A) consisted of an N-terminal red fluorescent protein tdTomato followed by the SspB micro and the full-length human FBP17 S isoform (tdTomato-micro-FBP17FL).^32^ The bait component of the optogenetic system was an iLID targeted to the plasma membrane with a CAAX prenylation sequence and tagged with the green fluorescent protein EGFP (iLID-EGFP-CAAX). Upon co-expression of the prey and the bait in COS7 African green monkey kidney cells, we found that iLID-EGFP-CAAX was mostly located on the plasma membrane while tdTom-micro-FBP17 was both on the plasma membrane and in the cytosol. In agreement with previous reports that overexpression of FBP17 results in the formation of positive membrane curvatures in the form of invaginating membrane tubules, we observed that COS7 cells overexpressing tdTom-micro-FBP17, with or without iLID-EGFP-CAAX, show some membrane tubulations in the absence of blue light (Supplementary Figure S1). Pulsed blue light (200 millisecond pulse duration at 5.3 W/cm^2^ on a fluorescence microscope) was delivered every 10 seconds both to image GFP-tagged constructs and to initiate iLID/SspB interactions. Blue light illumination resulted in a significant enhancement of the membrane tubules that are observable in both channels (Figure 2B, with full field images in Supplementary Figure S2). Additionally, we have switched the iLID and micro locations in the bait and prey constructs (tdTomato-iLID-FBP17FL + micro-EGFP-CAAX). The switched version shows similar behavior, i.e. some membrane tubules in the dark and significantly enhanced membrane tubules upon blue light stimulation (Supplementary Figure S3).

**Figure 2:**
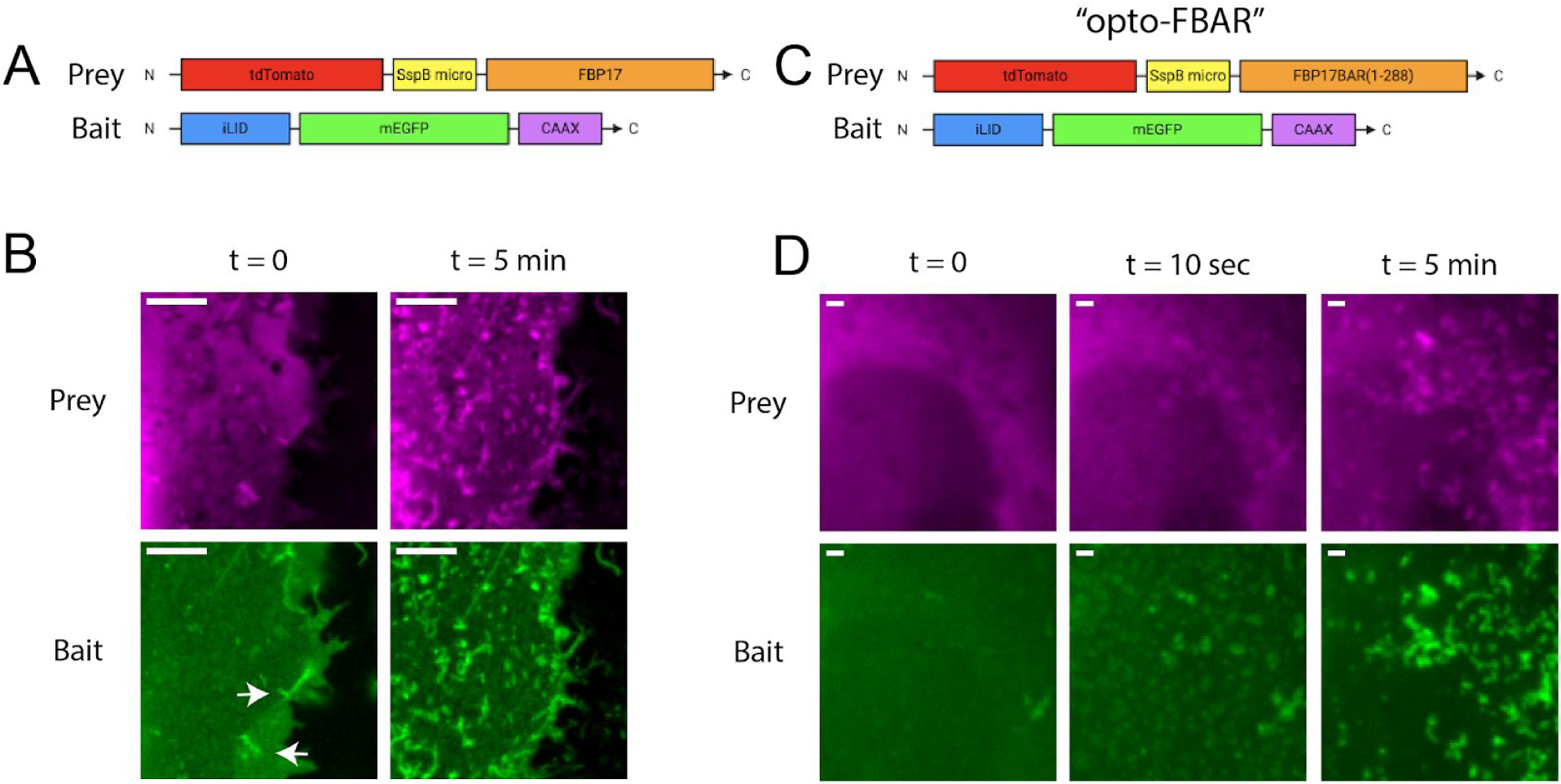
The construction and optimization of opto-FBAR. **A**) Box drawing of the initial design of the opto-FBAR system using full length FBP17. **B**) Zoomed images of COS7 cells co-expressing tdTom-SspB-FBP17 and iLID-EGFP-CAAX shown in A. In the dark (t=0), cells show some FBP17-positive membrane tubules (arrows). Blue light stimulation (t=5 min) drastically increased the amount of membrane tubules that is positive in both channels. Full field images available in Supplementary Figure S2. Scale bar = 5 μm. **C**) Box drawing of final opto-FBAR system using the FBAR domain, i.e. truncated FBP17(aa1-288). **D**) Zoomed-in images of COS7 cells expressing the final opto-FBAR system shown in C. tdTom-SspB-FBAR is completely cytosolic in the dark. Blue light rapidly recruits the prey to the membrane and result the formation of membrane tubules positive in both channels. Full field images available in Supplementary Figure S4. Scale bar = 1 μm.

To reduce the dark background, i.e. tubule formation without blue light stimulation, we chose to truncate FBP17 to include only the FBAR domain (aa 1-288) (Figure 2C). The FBAR domain of FBP17 is necessary and sufficient for curvature sensing.^26,33,34^ Upon transient transfection, we found that the truncated tdTom-micro-FBAR construct appeared to be almost exclusively cytosolic and did not result in the formation of membrane tubulation, whereas similar expression levels of full-length FBP17 constructs showed considerable amounts of tubulation before light. The observation that the FBP17 FBAR has a lower affinity for the membrane than full-length FBP17 has also been suggested in a recent study that shows multiple regions of FBP17 outside of its FBAR domain contributing to its membrane binding.^35^ Upon co-expression of tdTomato-micro-FBP17BAR and iLID-EGFP-CAAX, blue light elicited a robust membrane tubulation response (Figure 2D, with full field images in Supplementary Figure S4). Red channel images show tdTom-micro-FBP17BAR is highly cytosolic before addition of blue light. Membrane recruitment and tubule formation is apparent after 10 seconds of blue light exposure and significantly increases at 5 minutes. Green channel images of the iLID-EGFP-CAAX show colocalized tubulation formation, indicating that the FBP17 tubules formed by opto-FBAR are indeed curved features of the plasma membrane. We named the final tdTom-micro-FBP17BAR + iLID-EGFP-CAAX system “opto-FBAR.”

With multi-component protein systems such as opto-FBAR, varying transfection efficiencies between plasmids can result in large cell-to-cell variability in the ratio of their expression levels. To mitigate this, we chose to clone the CMV expression cassettes of both the bait and prey into a single plasmid to provide roughly equal expression levels of each component, as such has been shown with up to five expression cassettes.^36^ Single plasmids encoding both opto-FBAR components are used in the following studies for quantitative measurements.

### Reversible and subcellular generation of positive membrane curvature with opto-FBAR

With opto-FBAR we are able to reversibly generate positive membrane curvature with a blue light stimulus. As shown in Figure 3A, blue light stimulation for 5 min induced dramatic formation of membrane tubules. About 30 min after blue light was turned off, these membrane tubules had mostly disappeared and tdTom-micro-FBP17BAR became cytosolic (full field images in Supplementary Figure S5). Opto-FBAR affords subcellular generation of positive membrane curvature owing to the relatively slow diffusion of iLID-EGFP-CAAX tethered on the plasma membrane and the fast dissociation kinetics of iLID and micro heterodimer in the dark.^21^ As shown in Figure 3B (full field images in Supplementary Figure S6), blue light focused on a small region of the plasma membrane elicited the formation of membrane tubules only in the illuminated area.

**Figure 3:**
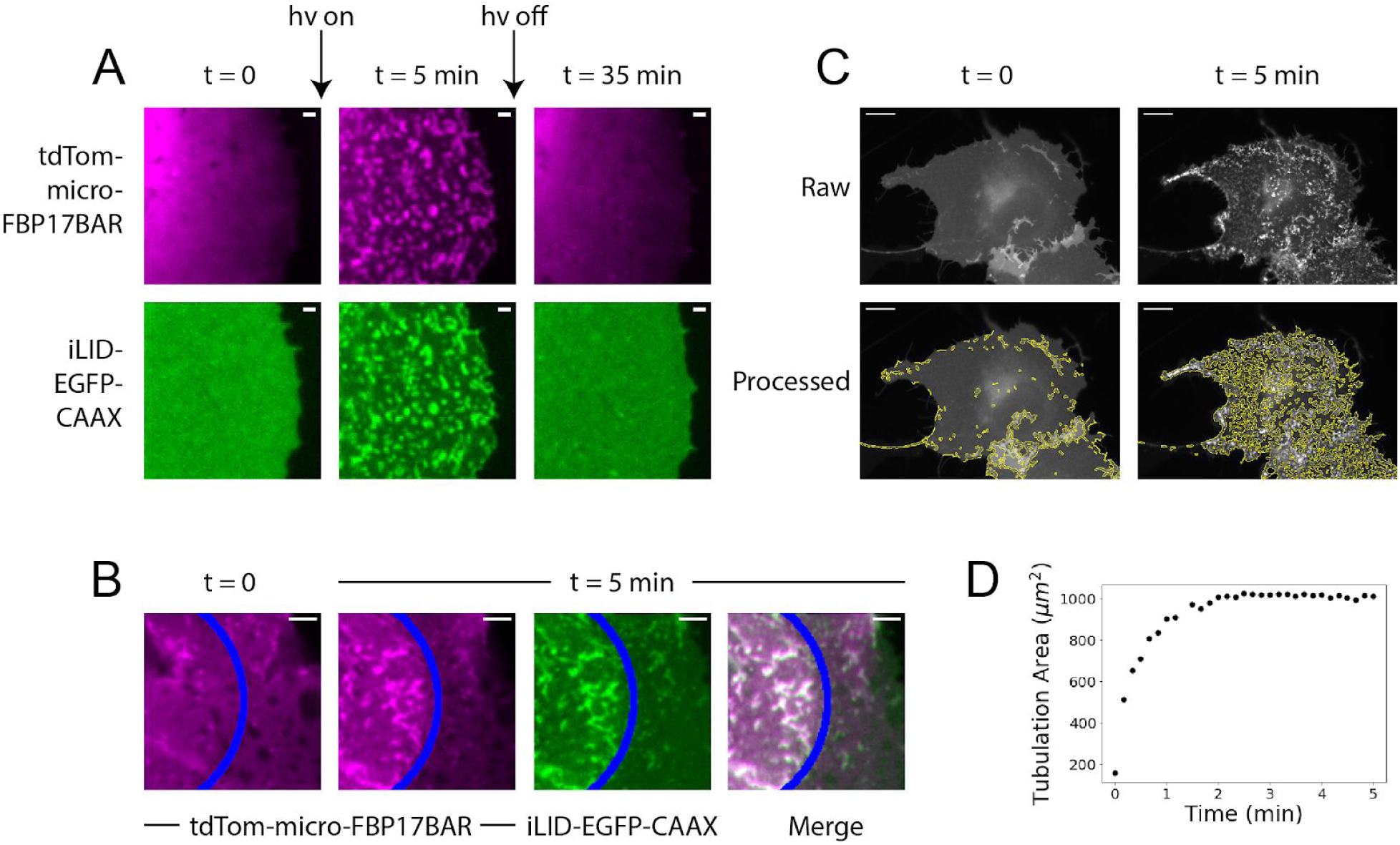
Spatiotemporal controlled generation of positive membrane curvature by opto-FBAR. **A**) COS7 cells expressing the opto-FBAR system were exposed to blue light for 5 minutes, which induced significant membrane tubules. Light-induced positive membrane curvatures are fully reversed after 30 min in the dark. Full field images available in Supplementary Figure S5. Scale bar = 1 μm. **B**) A subcellular region (areas to the left of the blue line) of COS7 cells expressing opto-FBAR system were exposed to blue light for 5 minutes. Membrane tubules are induced in the region exposed to blue light. Full field images available in Supplementary Figure S6. Scale bar = 5 μm. **C**) Green channel (iLID-EGFP-CAAX) were analyzed with TubuleTracer to quantify the tubulation extent over time. The results are shown under the raw images. Scale bar = 10 μm. **D**) The plot of tubulation area vs time from the cell shown in C. Tubulation reaches a maximum around 2 min.

We quantified the kinetics of the curvature generated by opto-FBAR by automated data analysis. Recently, a freely available ImageJ macro AxonTracer was published with the intended use of identifying fluorescently tagged axons in tissue sections using Gaussian-scaled derivatives, providing an output measurement of total axon length.^37^ Given that the relative scaling of an axon to brain tissue is similar to that of a membrane tubule to a cell, we used a modified version of AxonTracer to identify membrane tubules formed by opto-FBAR with reasonable accuracy (Figure 3C, see method section for details). This program, which we named “TubuleTracer,” shows that the total length of membrane tubules rapidly increases with respect to the duration of blue light stimulation and reaches a maximum after approximately 2 min (Figure 3D).

### Engineering an opto-IBAR system for negative curvature generation

We next sought to develop an optogenetic system that generates negative membrane curvature using the inverse BAR protein IRSp53.^28^ We first compared the expression of full length IRSp53 and IBAR (a.a. 1-300) for inducing negative membrane curvature, i.e. membrane protrusions. Unlike the FBAR domain that is cytosolic and inactive, we found that IBAR-mEGFP appeared to be mostly membrane bound and causes more dramatic membrane protrusions than full length IRSp53-mEGFP (Figure 4A, full images in Suppl. Figure S7). This result shows that it is not feasible to use the same and simple opto-FBAR approach to construct opto-IBAR.

**Figure 4:**
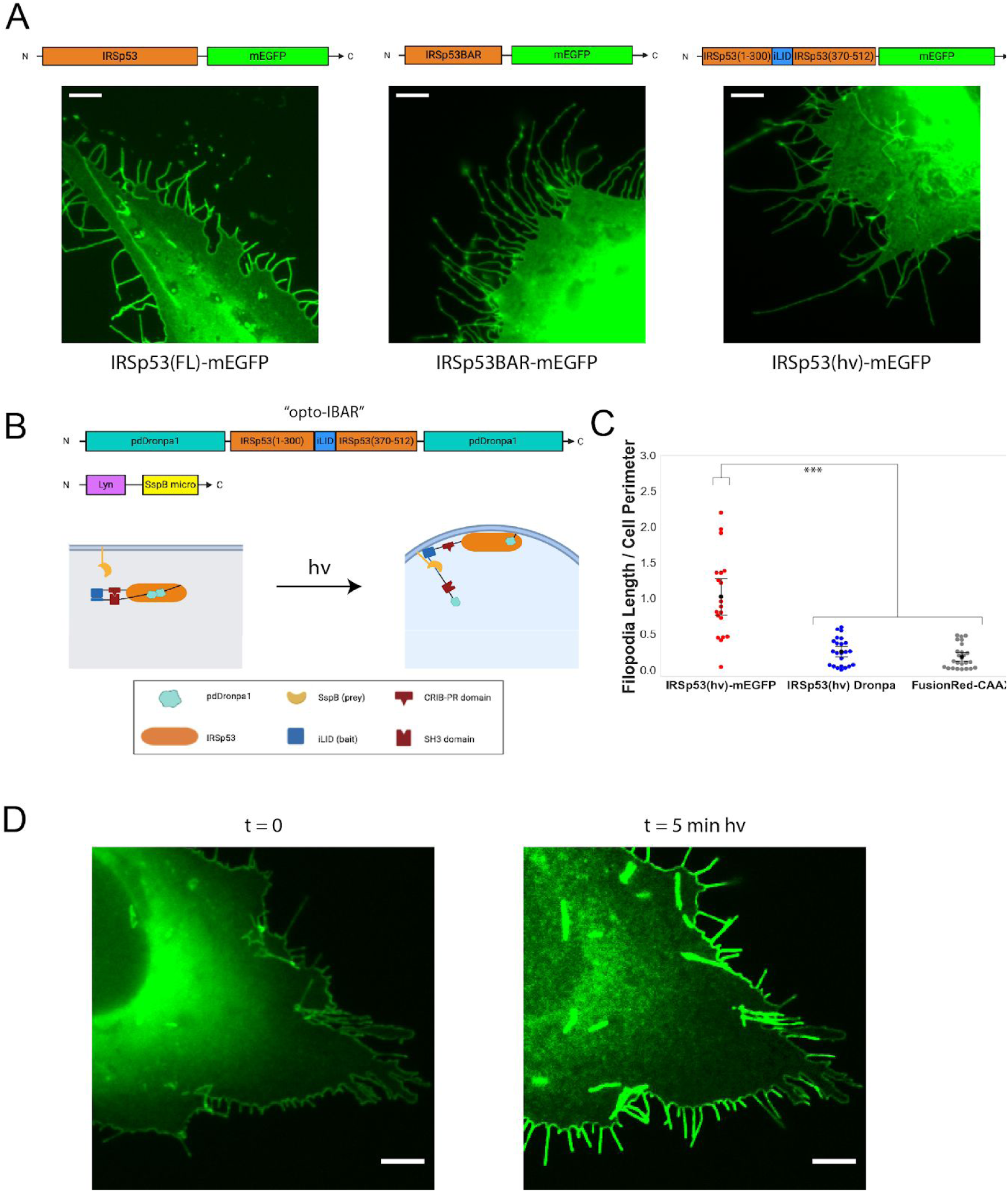
The construction and optimization of opto-IBAR. **A**) U2OS cells overexpressing indicated IRSp53 constructs show substantial amount of filopodia protrusions. Scale bars = 5 μm. Full field images available in Supplementary Figure S7. **B**) Schematic depicting IRSp53 photocaging by pdDronpa1 and the final opto-IBAR constructs. **C**) Swarm plots of total filopodia length normalized to cell perimeter. ***p < 0.005. **D**) Images of U2OS cell expressing opto-IBAR construct before (left) and after (right) 5 minutes of blue light activation show dramatic increase of the filopodia length. Scale bars = 5 μm. *Note:* Images were bleach corrected using the histogram matching method in ImageJ. Full field images available in Supplementary Figure S8.

It has been shown that IRSp53 exhibits an autoinhibitory regulation mechanism at its CRIB-PR domain that can be activated by Cdc42 binding.^38^ In addition, this autoinhibition is regulated via phosphorylation and 14-3-3 binding at residues T340, T360, and S366 between the CRIB-PR and SH3 domains.^39^ We desired an optogenetic system that is not affected by such regulation, thus we replaced amino acids 300-370 of the IRSp53 S isoform with iLID, removing the 14-3-3 binding sites but maintaining the other known functional domains. We herein refer to this IRSp53 construct as IRSp53(hv). Expression of IRSp53(hv)-mEGFP alone induces significant membrane protrusions similar to that by IRSp53-mEGFP (Figure 4A, **right**). This high dark background makes IRSp53(hv)-mEGFP unsuitable for constructing opto-IBAR.

For optogenetic control, we desire an optogenetic construct that has low membrane affinity without light, but retains the ability to oligomerize after blue light recruit it to the membrane. In order to reduce the membrane affinity of the IRSp53(hv)-mEGFP protein, we sought a caging system by using photodissociable dimeric Dronpa (pdDronpa). In the dark, two intra-molecular pdDronpa domains interact with each other to form a homodimer, which can cage the protein domain of interest in between. Exposure of blue/cyan light results in dissociation of the pdDronpa dimer and therefore releases the caged protein of interest. This approach was utilized to develop single-chain photoswitchable kinases which operate via steric occlusion of the kinase active site, as well as a photoswitchable CRISPR-Cas9 architecture for light-dependent gene interactions.^40,41^

We designed the caged IRSp53 by fusing pdDronpa1 domains to the N-terminus and the C-terminus of IRSp53(hv) (Figure 4B). In the dark, the two N- and C-pdDronpa1 domains will interact to form a homodimer to inhibit IRSp53, and blue light exposure would simultaneously result in dissociation of the pdDronpa1 homodimer and the association of iLID and micro for recruitment to the plasma membrane. We constructed a pdDronpa1-IRSp53(hv)-pdDronpa1 plasmid with a flexible 3xGSS linker between the N-terminal pdDronpa1 and IRSp53(hv), and a 2xGSS linker between the IRSp53(hv) and C-terminal pdDronpa1. We hereafter refer to this construct as IRSp53(hv) Dronpa. Before blue light stimulation, the filopodia level (dark background) of IRSp53(hv) Dronpa is comparable or slightly higher than the control system expressing FusionRed-CAAX (Figure 4C). We cloned the Lyn-micro expression cassette into a plasmid encoding IRSp53(hv) Dronpa to limit variabilities in the bait and prey expression ratio, as discussed earlier for opto-FBAR. We chose a Lyn membrane targeting signal because it is believed to target membrane rafts, which are known to be concentrated at filopodia.^42^ When IRSp53(hv) Dronpa was coexpressed with Lyn-micro, we found this optogenetic system gave strong and robust responses of filopodia formation. Five minutes of blue light stimulation induced a dramatic increase in the filopodia number, the average length of filopodia, and the total filopodia length (Figure 4E). We chose to conclude our engineering efforts with the IRSp53(hv) Dronpa and Lyn-micro constructs, designating them as “opto-IBAR.”

### Characterization of the negative curvature generation with opto-IBAR

By performing FiloQuant analysis on live cells expressing opto-IBAR, we are able to measure how the length and number of filopodia change over time. As shown in Figure 5A, cells are devoid of long filopodia prior to light stimulation. Upon blue light exposure, opto-IBAR is quickly activated that induced both the formation of new filopodia and the elongation of existing protrusions at the cell edge. Kinetic analysis shows that the filopodia formation reaches a maximum around 5 min (Figure 5A).

**Figure 5:**
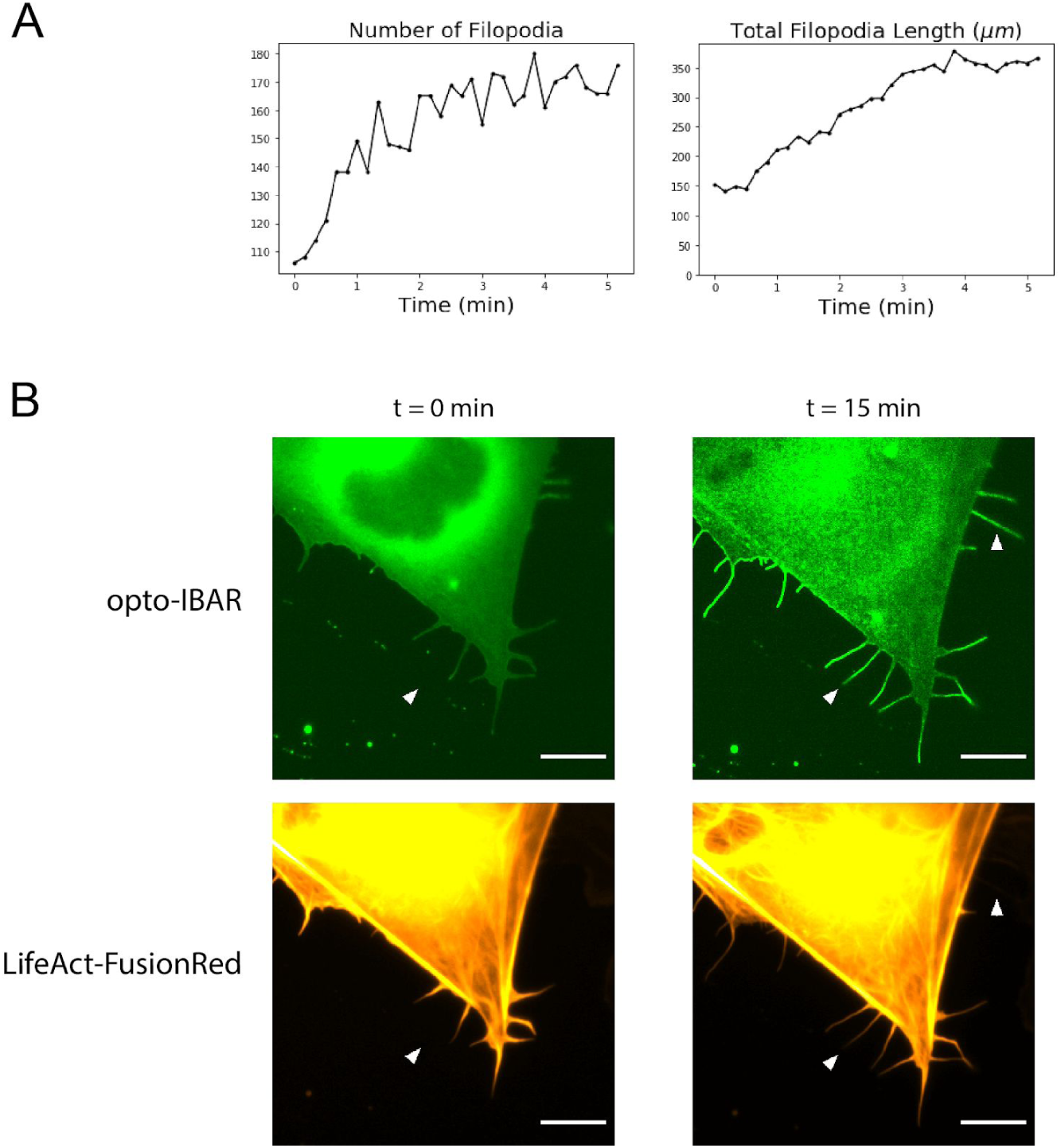
**A**) Plots of filopodia number and total filopodia length from FiloQuant analysis of cell shown in Figure 4D. **B**) Zoomed images of U2OS cells expressing the opto-IBAR system and a LifeAct-FusionRed F-actin marker. Cells were activated with blue light for 15 minutes as described in Methods. Most filopodia contain F-actin, but some long filopodia are devoid of F-actin (arrows). Scale bars = 5 μm. *Note:* Green channel images were bleach corrected using the histogram matching method in ImageJ. Full field images available in Supplementary Figure S9.

Our opto-IBAR construct contains the IRSp53 SH3 domain that is capable of interacting with the actin cytoskeleton. When co-expressing opto-IBAR with LifeAct-FusionRed in U2OS cells, we found that most filopodia induced by opto-IBAR activation are indeed also positive for actin fibers (Figure 5B, **lower arrow**). However, we found that some filopodia are devoid of actin, suggesting that opto-IBAR polymerization on the membrane is sufficient to drive the formation of filopodia (Figure 5B, **upper arrow**).

Here, we have developed strategies that achieve light-inducible generation of either positive or negative membrane curvatures in live cells using BAR domain proteins. The key aspect of our strategies is identifying a protein construct that has a low membrane affinity, yet is able to oligomerize and generate membrane curvature when forcibly recruited to the membrane. For opto-FBAR, the FBAR domain protein is cytosolic and its membrane recruitment by blue light induces the formation of positive membrane curvature, with high enough spatiotemporal precision to induce membrane curvature at the subcellular level. In our efforts to engineer opto-IBAR, we incorporated a second optogenetic system pdDronpa. Opto-IBAR has minimal membrane binding but can be recruited to the membrane by blue light and induce negative membrane protrusions. This approach for controlling membrane curvature is simple and effective, and is likely to be expandable to other BAR domain proteins.

## Methods

### Plasmid construction

pLL7.0: Venus-iLID-CAAX (Addgene plasmid # 60411) and pLL7.0: tgRFPt-SSPB WT (Addgene plasmid # 60415) were gifts from Brian Kuhlman.^19^ The pdDronpa1 sequence was obtained from a synthesized gblock (Integrated DNA Technologies). All plasmid constructs were cloned using Gibson Assembly.^43^

### Cell culture and transfection

U2OS cells (ATCC® HTB-96™) and COS-7 cells (ATCC® CRL-1651™) were maintained in complete medium (DMEM supplemented with 10% fetal bovine serum, both from Thermo Fisher Scientific) and grown in a standard incubator at 37°C with 5% CO2. Cells tested negative for mycoplasma contamination by PCR with primers 5’-GTGGGGAGCAAAYAGGATTAGA-3’ and 5’-GGCATGATGATTTGACGTCRT-3’.^44^

COS-7 cells were plated on poly-L-lysine coated coverslips and transfected using Lipofectamine 2000 (Thermo Fisher Scientific) according to the manufacturer’s protocol. The transfected cells were allowed to recover overnight in complete culture medium. U2OS cells were transfected via electroporation using the Amaxa Nucleofector II (Lonza). Cells were added to a suspension of DNA in electroporation buffer (7 mM ATP, 11.7 mM MgCl_2_, 86 mM KH_2_PO_4_, 13.7 mM NaHCO_3_, 1.9 mM glucose), transferred to a 2mm electroporation cuvette (Fisher Scientific) and subjected to the manufacturer provided protocol for U2OS cells, then plated on coverslips that were pre-coated with a matrix consisting of 0.02% gelatin from porcine skin (Sigma-Aldrich G1890) and 4 ug/mL Fibronectin bovine plasma (Sigma-Aldrich F1141). Cells were allowed to recover overnight in complete medium.

### Live cell imaging

Prior to imaging, the culture medium was changed to an imaging medium derived from ref. 23 containing increased glucose to prevent starvation-induced filopodia:^45^ 20 mM HEPES at pH 7.5, 150 mM NaCl, 5 mM KCl, 1 mM CaCl2, 1 mM MgCl2 and 4.5 g/L glucose supplemented with 10% fetal bovine serum. Live cell imaging was performed on an epifluorescence microscope (Leica DMI6000B) equipped with an on-stage CO2 incubation chamber (Tokai Hit GM-8000) and a motorized stage (Prior). An adaptive focus control was used to actively keep the image in focus during the period of imaging. A light-emitting diode (LED) light engine (Lumencor) was used as the light source for fluorescence imaging. Pulsed blue light (200 ms pulse duration at 5.3 W/cm^2^) was delivered every 10 s both to image GFP-tagged constructs and to initiate iLID/SspB interactions. Pulsed green light (200 ms pulse duration) was used to image mCherry or tdTomato. The microscope was equipped with a commercial GFP filter cube (Leica; excitation filter 472/30, dichroic mirror 495, emission filter 520/35) and a commercial Texas red filter cube (Leica; excitation filter 560/40, dichroic mirror 595, emission filter 645/75). All Images were acquired with an oil-immersion 100× objective. All experiments were imaged with a sensitive CMOS camera (PCO.EDGE 5.5) (PCO).

### Filopodia assay

Transfected U2OS cells were fixed and visualized using fluorescence microscopy on the same Leica system described, without the on-stage CO2 incubation chamber. To prevent the loss of filopodia from standard fixation methods, the MEM-fix fixation protocol was used with two modifications.^46^ First, cells were not re-seeded after transfection, and second, as permeabilization of the cells was not necessary post-fixation, cells were instead treated with 1 mg/mL sodium borohydride alone in pure water for 1-2 hrs to reduce autofluorescence.

### Development of TubuleTracer

We modified the freely available ImageJ macro AxonTracer to identify membrane tubules formed by opto-FBAR. First, we removed the region-of-interest identification steps, as plasma membrane positive curvature will always be inside of the cell area. After optimization of parameters such as hole identification settings and the Gaussian smoothing factor, the initial program was able to identify membrane tubulation with reasonable accuracy. In the case of clustered tubules, the program’s skeletonization step of the identified tubulation area had a tendency to reduce the clusters to single tubules. Thus we removed the skeletonization step, altering the output measurement from total tubulation length to tubulation area. This program, which we named “TubuleTracer,” showed a marked improvement in the ability to quantify tubule clusters and is available in the Supplementary Information (Supplementary File 1).

## Supporting information

Supplementary Video 1: COS7 cell expressing opto-FBAR system. Time = min:sec

Supplementary Video 2: U2OS cell expressing opto-IBAR system. Video bleach corrected using the histogram matching method in ImageJ. Time = min:sec

Supplementary Video 3: Supplementary Video 2 without bleach correction. Time = min:sec

## Acknowledgements

This work was supported by the US NIH (<u>1R01GM128142</u>). All schematics were created using BioRender.com. Electrostatic surface potential maps were generated using PyMol.

---

**Supplementary Figure S1:**
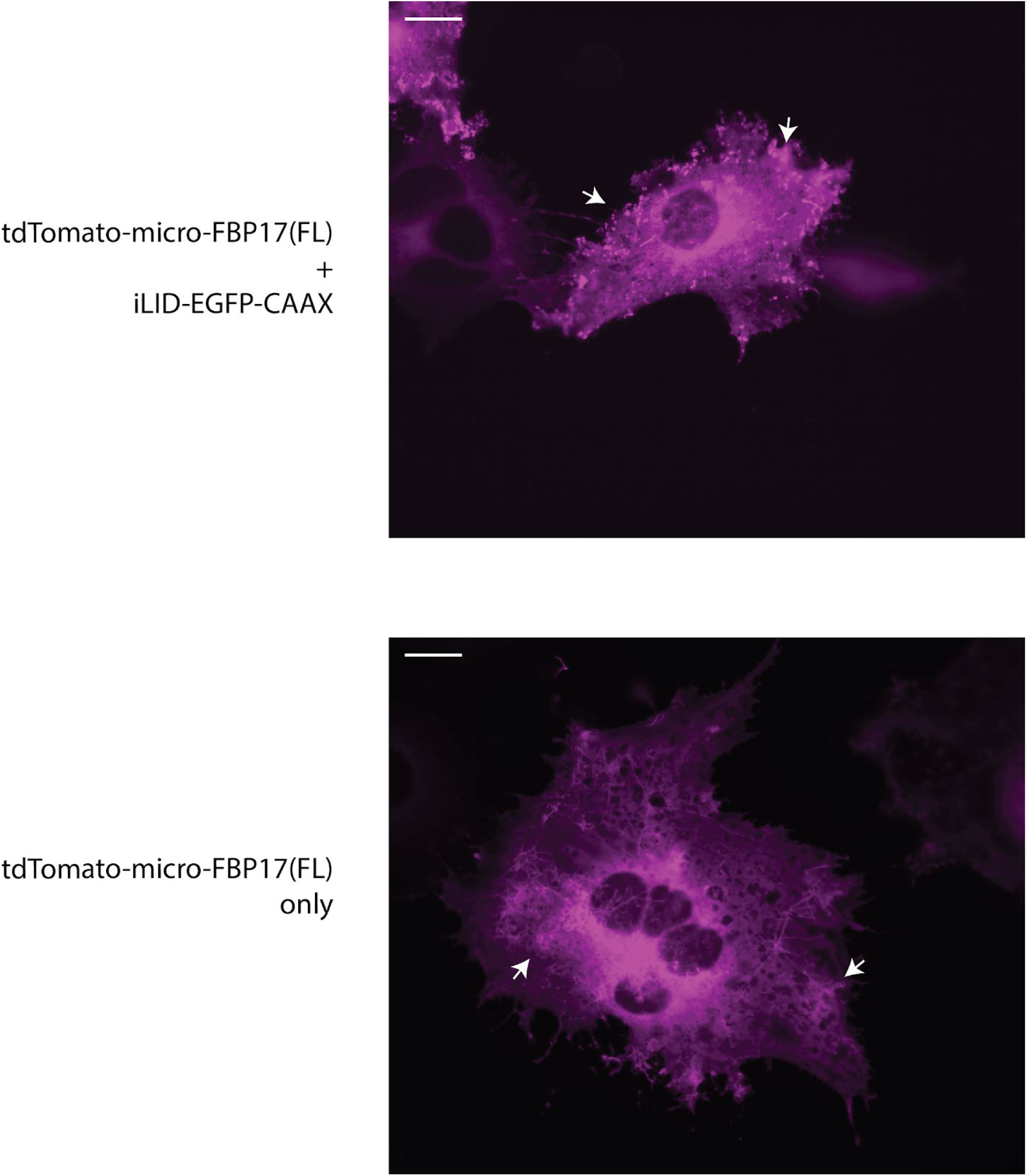
opto-FBAR with FBP17(FL) exhibits high levels of tubulation (arrows) in the dark. *Top:* COS7 cells expressing tdTom-micro-FBP17(FL) with iLID-EGFP-CAAX in the dark. *Bottom*: COS7 cells expressing tdTom-micro-FBP17(FL) only. Images show the red channel (tdTom-micro-FBP17(FL)). Scale bar = 10 μm.

**Supplementary Figure S2:**
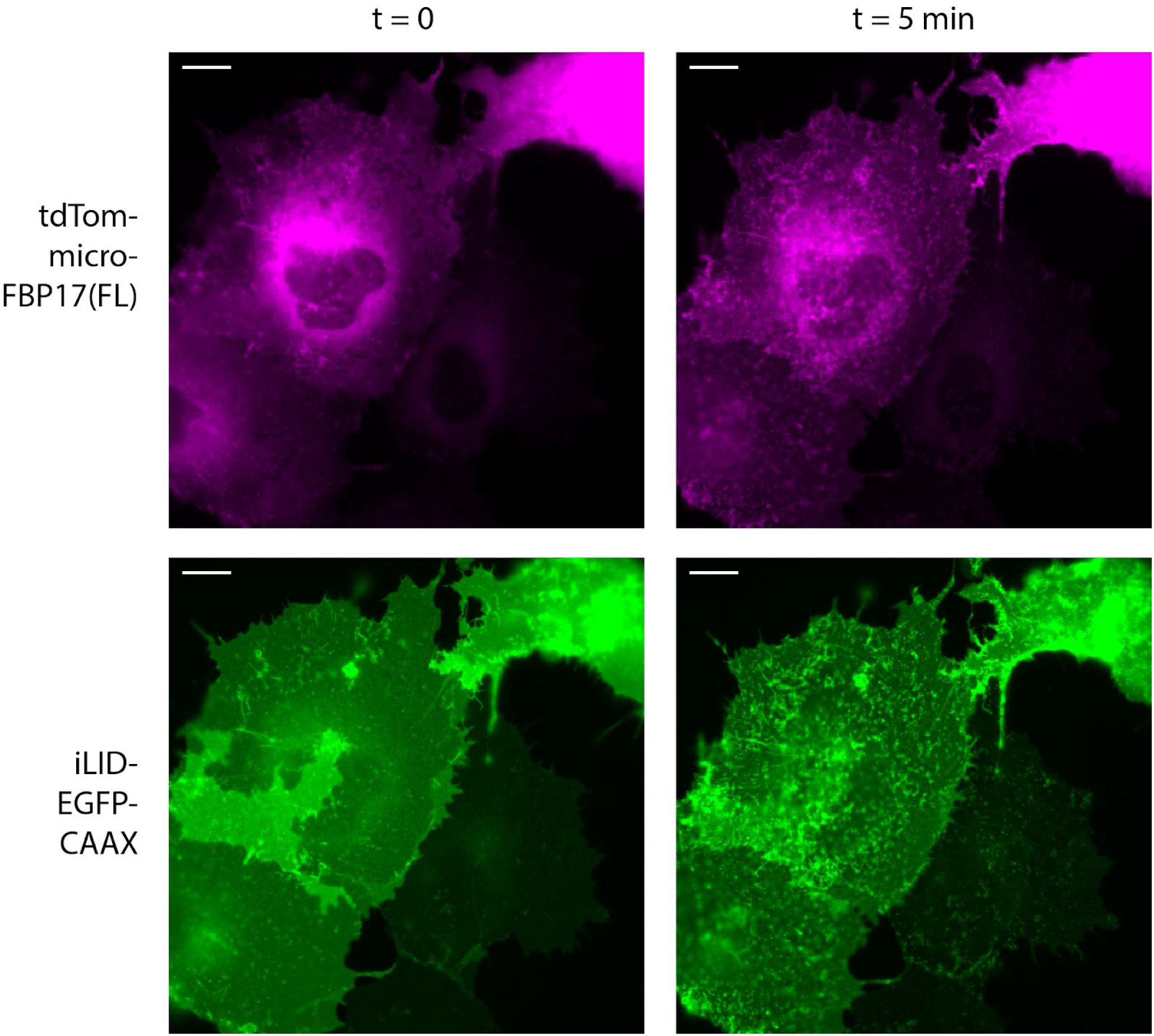
Full field images of cells shown in Figure 2B. Scale bar = 10 μm.

**Supplementary Figure S3:**
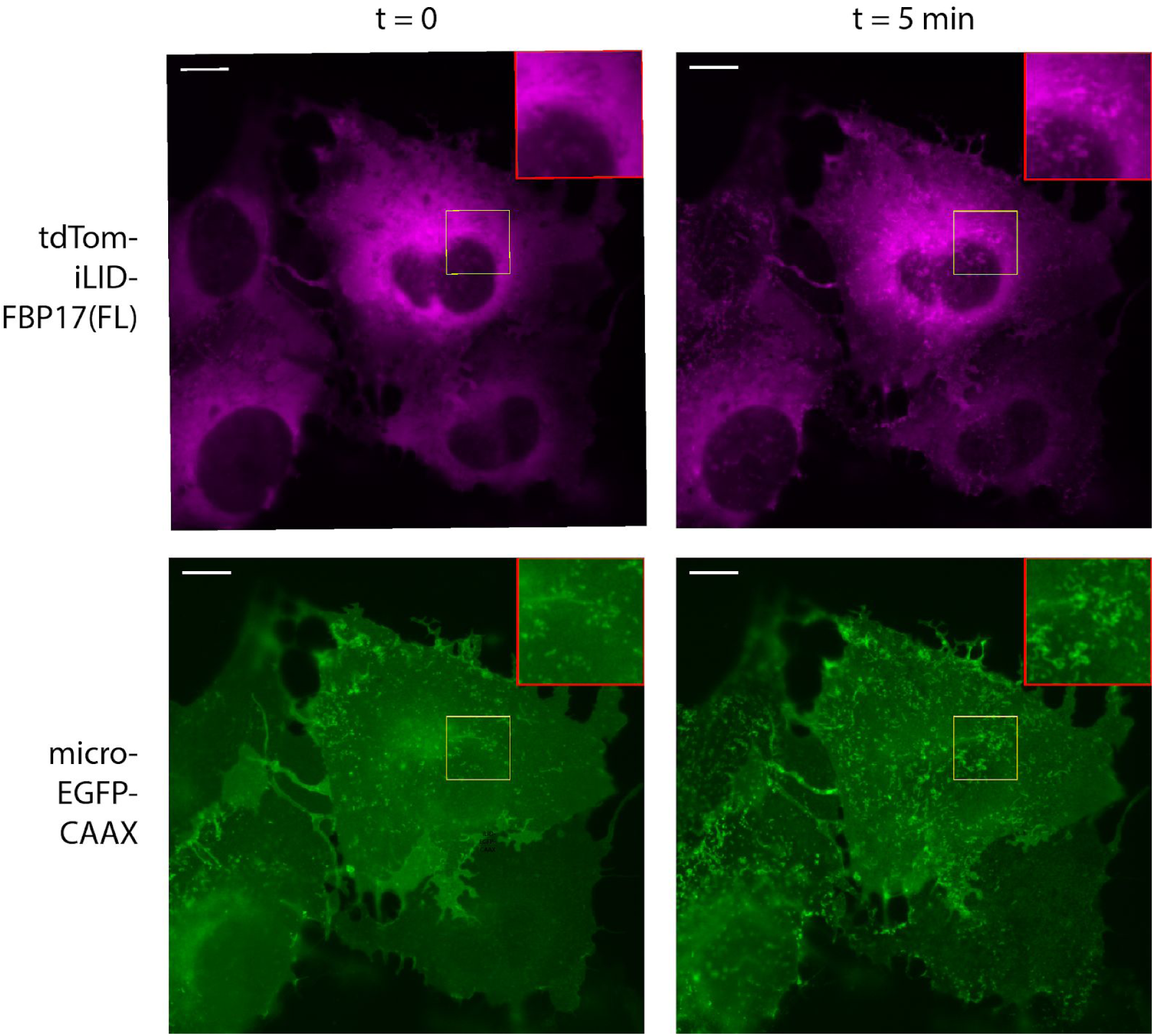
COS7 cells expressing tdtom-iLID-FBP17(FL) and micro-EGFP-CAAX, i.e. the bait and prey positions are switched. Cells were kept in the dark prior to 5 minutes of blue light exposure. Scale bar = 10 μm.

**Supplementary Figure S4:**
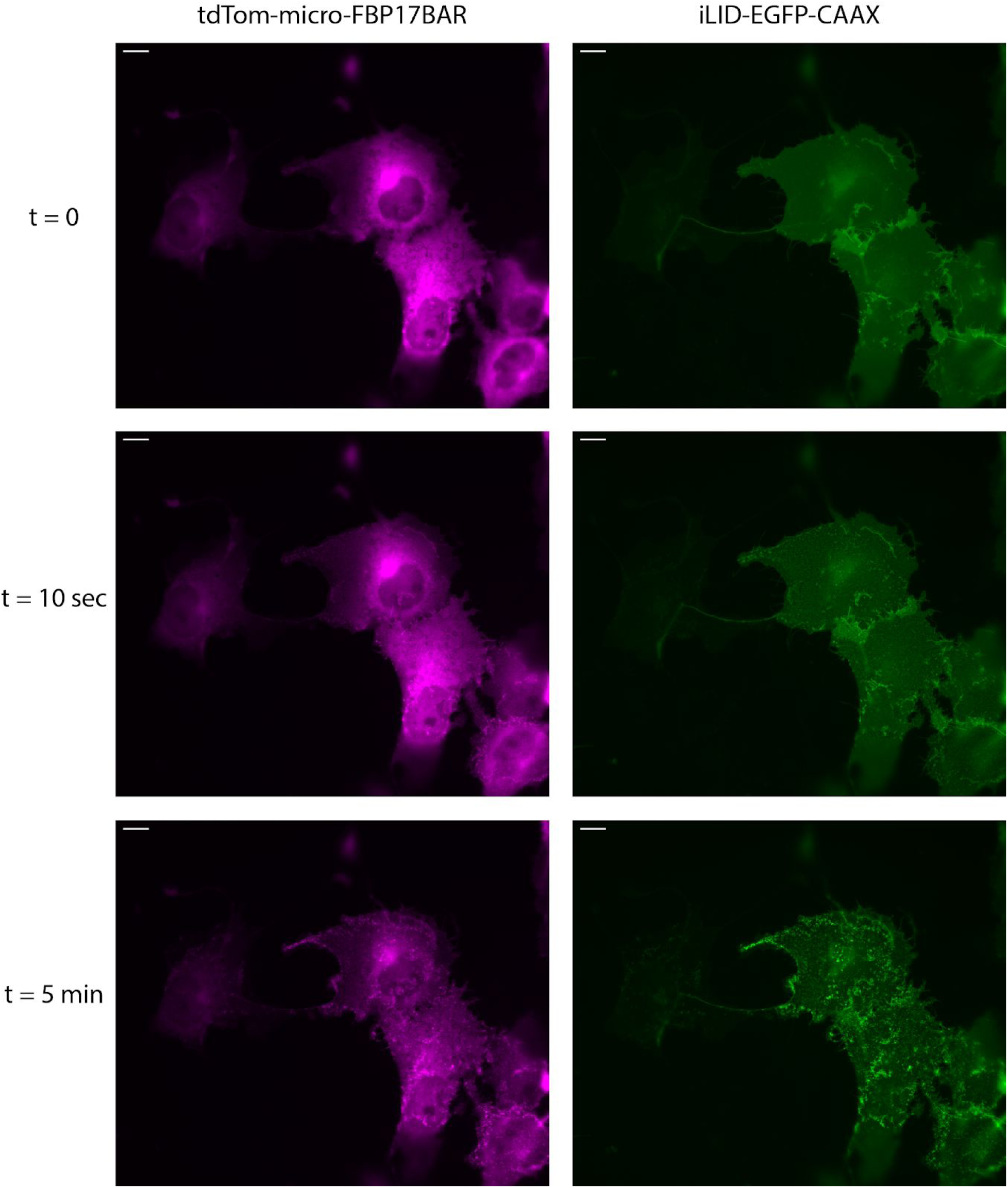
Full field images of cells shown in Figure 2D. Scale bar = 10 μm.

**Supplementary Figure S5:**
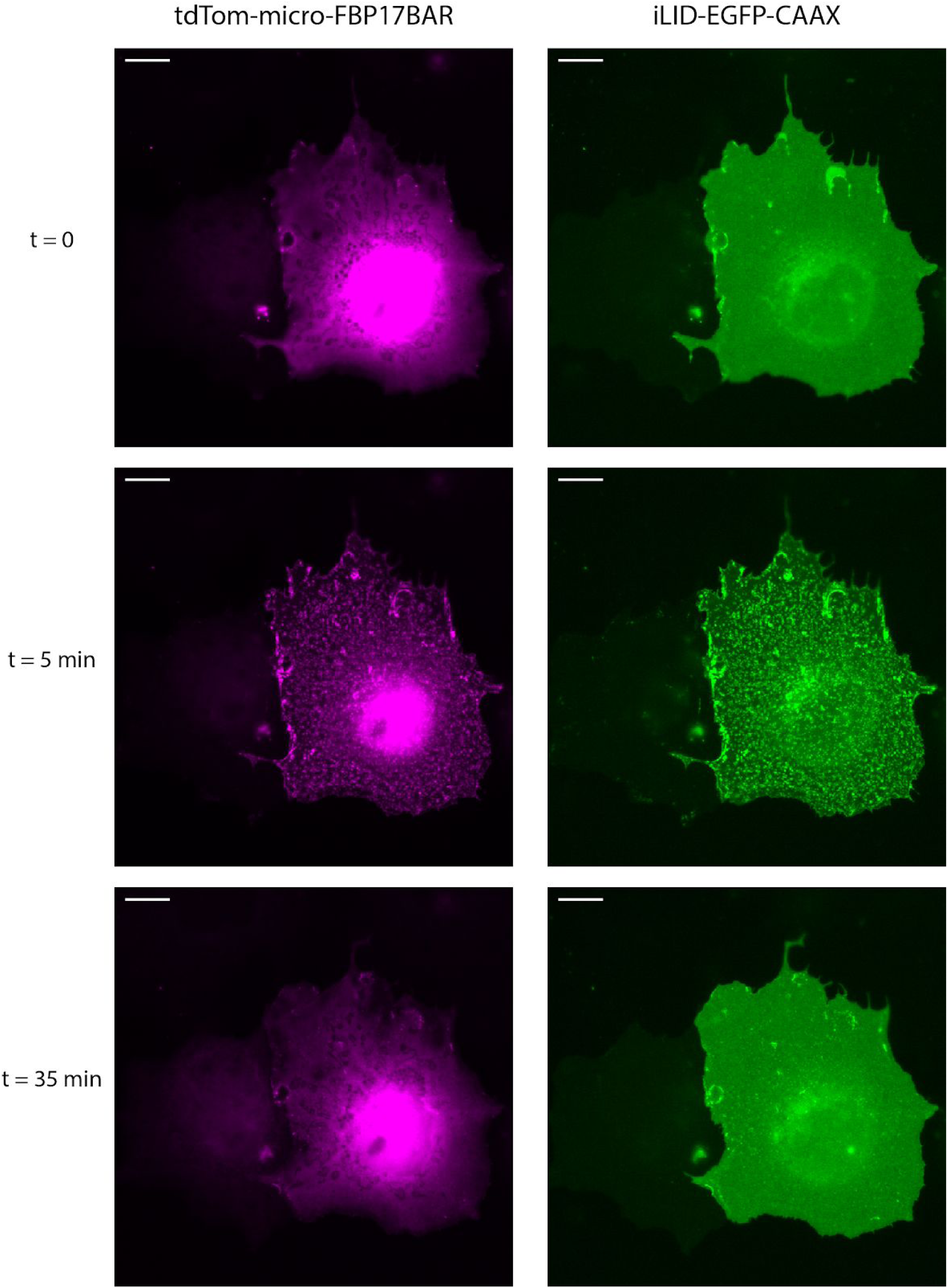
Full field images of cells shown in Figure 3A. Scale bar = 10 μm.

**Supplementary Figure S6:**
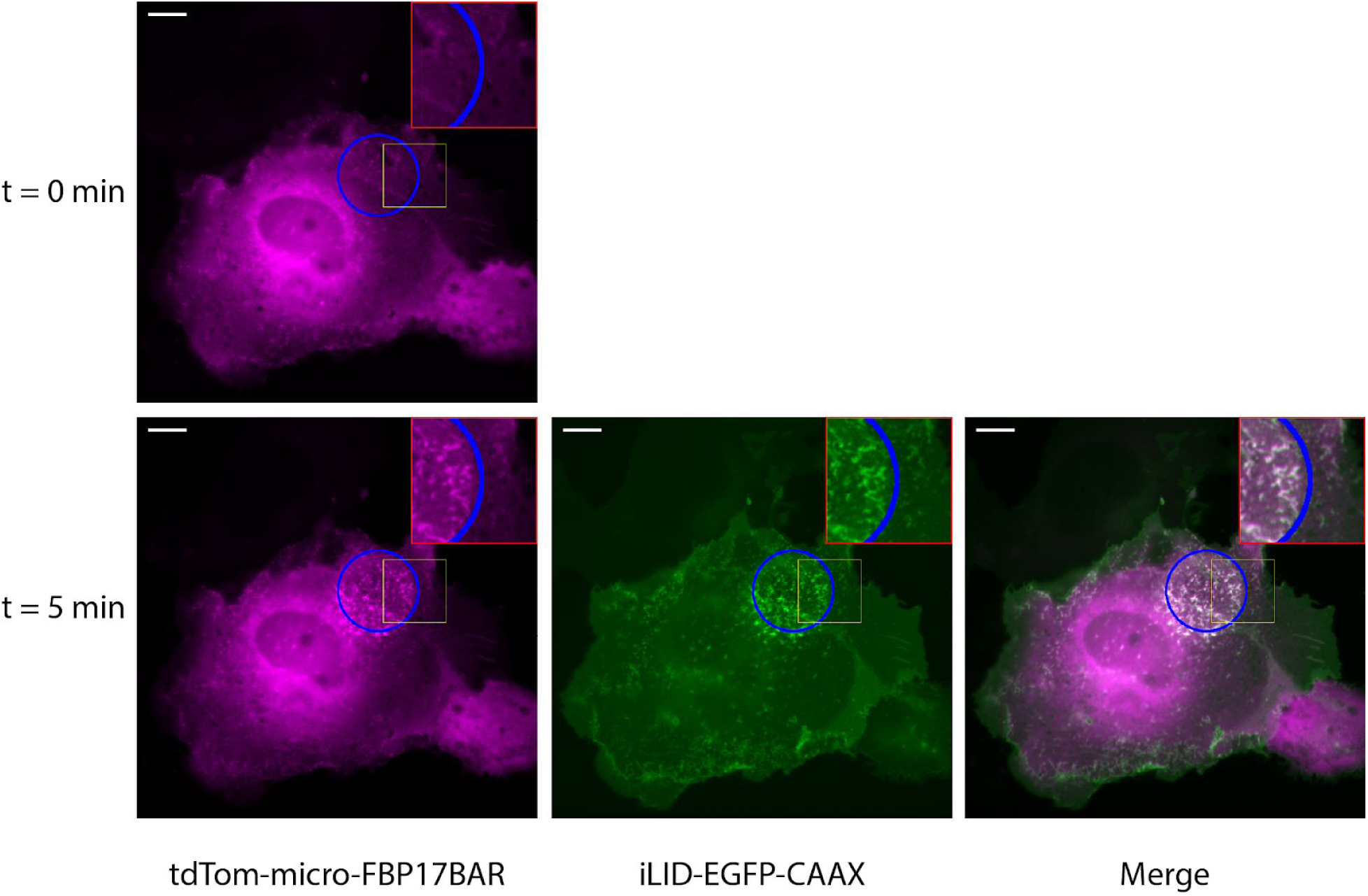
Full field images of cells shown in Figure 3B. Blue light illumination for 5 minutes was limited to a small area of the cell (blue circle). Yellow box indicates the area shown in Figure 3B. Scale bar = 10 μm.

**Supplementary Figure S7:**
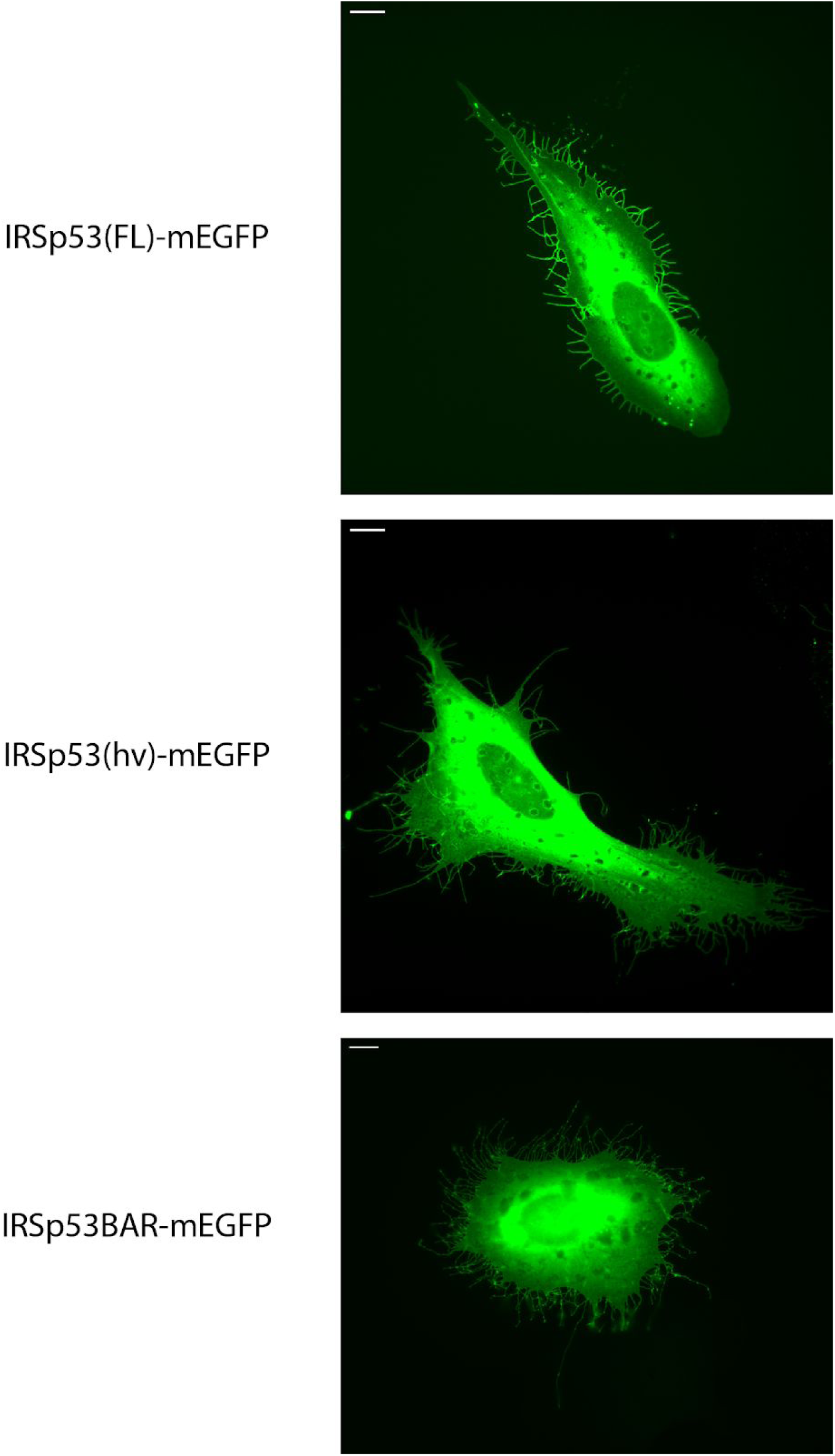
Full field images of cells shown in Figure 4A. Scale bar = 10 μm.

**Supplementary Figure S8:**
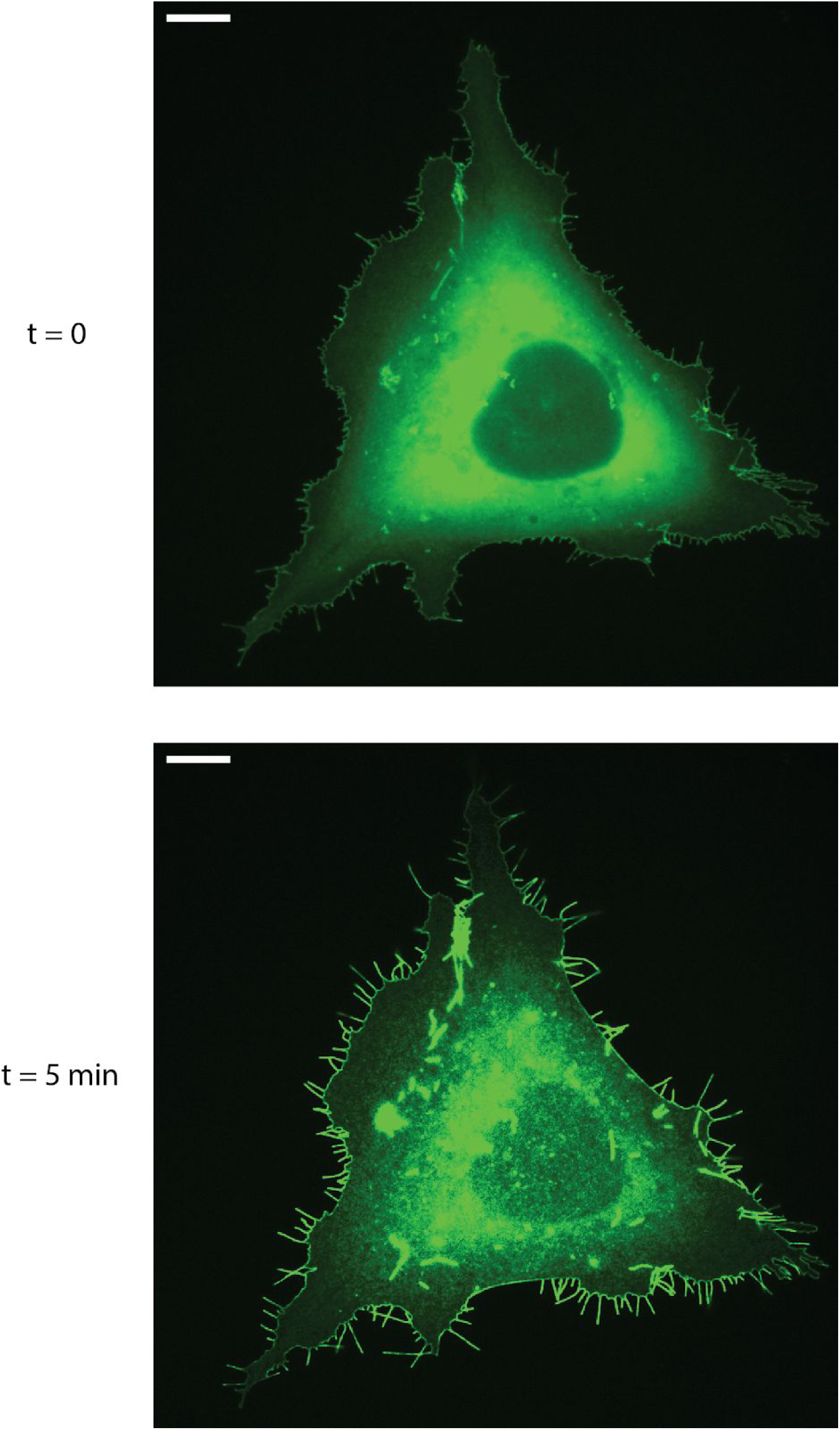
Full field images of cells shown in Figure 4D. Scale bar = 10 μm. *Note*: Images were bleach corrected using the histogram matching method in ImageJ.

**Supplementary Figure S9:**
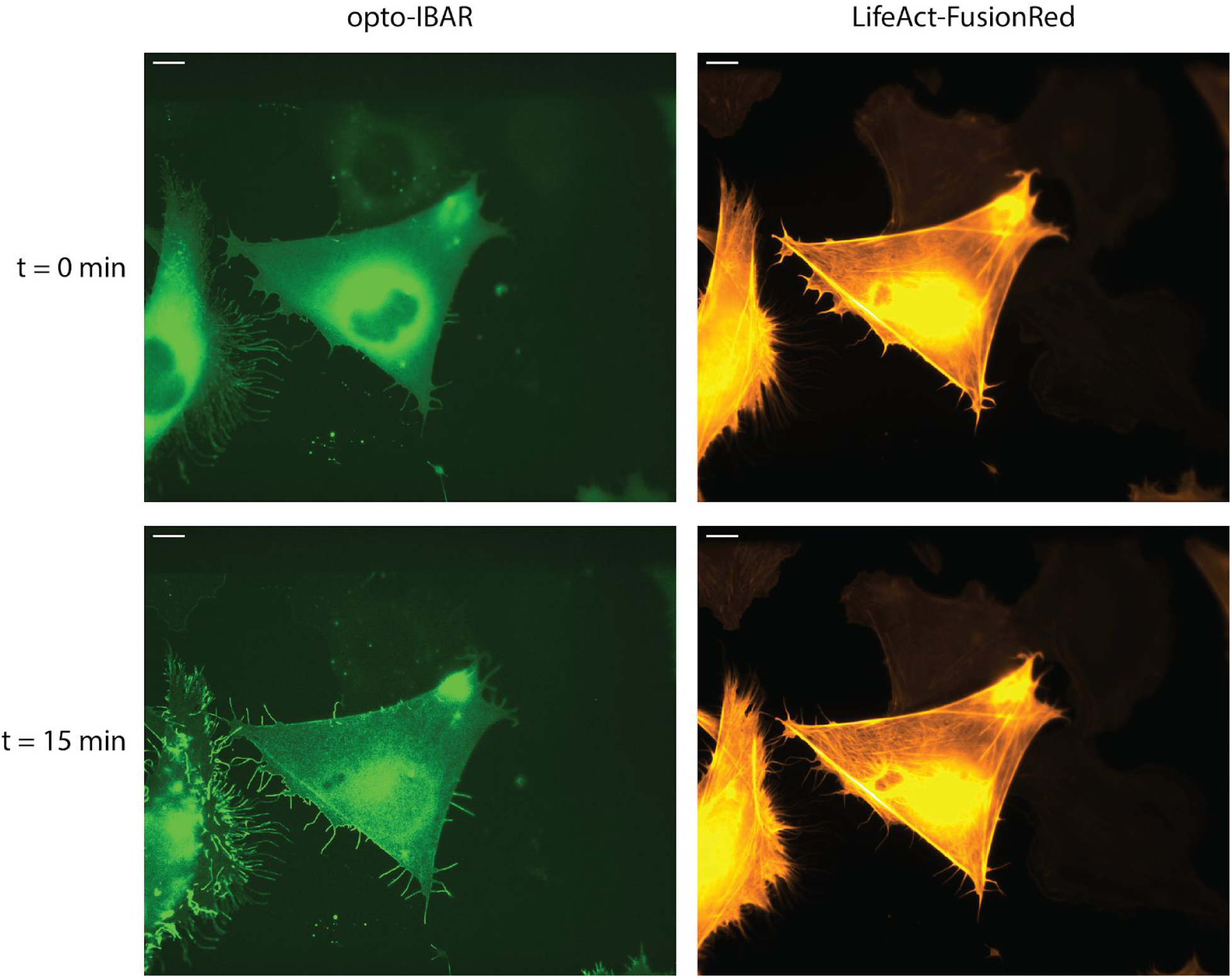
Full field images of cells shown in Figure 5B. Scale bar = 10 μm. *Note*: Green channel Images were bleach corrected using the histogram matching method in ImageJ.

